# Pixelwise H-score: a novel digital image analysis-based metric to quantify membrane biomarker expression from immunohistochemistry images

**DOI:** 10.1101/2021.01.06.425539

**Authors:** Sripad Ram, Pamela Vizcarra, Pamela Whalen, Shibing Deng, CL Painter, Amy Jackson-Fisher, Steven Pirie-Shepherd, Xiaoling Xia, Eric L. Powell

## Abstract

Immunohistochemistry (IHC) assays play a central role in evaluating biomarker expression in tissue sections for diagnostic and research applications. Manual scoring of IHC images, which is the current standard of practice, is known to have several shortcomings in terms of reproducibility and scalability to large scale studies. Here, by using a digital image analysis-based approach, we introduce a new metric called the pixelwise H-score (pix H-score) that quantifies biomarker expression from whole-slide scanned IHC images. The pix H-score is an unsupervised algorithm that only requires the specification of intensity thresholds for the biomarker and the nuclear-counterstain channels. We present the detailed implementation of the pix H-score in two different whole-slide image analysis software packages Visiopharm and HALO. We consider three biomarkers P-cadherin, PD-L1, and 5T4, and show how the pix H-score exhibits tight concordance to multiple orthogonal measurements of biomarker abundance such as the biomarker mRNA transcript and the pathologist H-score. We also compare the pix H-score to existing automated image analysis algorithms and demonstrate that the pix H-score provides either comparable or significantly better performance over these methodologies. We also present results of an empirical resampling approach to assess the performance of the pix H-score in estimating biomarker abundance from select regions within the tumor tissue relative to the whole tumor resection. We anticipate that the new metric will be broadly applicable to quantify biomarker expression from a wide variety of IHC images. Moreover, these results underscore the benefit of digital image analysis-based approaches which offer an objective, reproducible, and highly scalable strategy to quantitatively analyze IHC images.

## INTRODUCTION

Immunohistochemistry (IHC) is a core technology that is used to evaluate the spatial distribution and abundance of biomarkers at the protein level in tissue samples. In oncology clinical diagnosis and research applications, IHC assays play a central role in tumor characterization and biomarker assessment. Typically, IHC images are qualitatively evaluated by a trained expert, such as a pathologist, and in some cases this is complemented by a semi-quantitative score [1]. However, visual quantitative scoring of IHC images is not routinely performed due to several shortcomings. On the one hand, visual quantitative scoring is time consuming and is often not feasible to perform on a routine basis especially for large studies. On the other hand, visual quantitative scores are subjective and often have a limited dynamic range due to their categorical nature (e.g. manual scores of 0, 1+, 2+, and 3+). Consequently, they may not have the granularity to adequately capture biomarker expression from an IHC slide [2, 3]. The subjectivity of the scoring process, in turn, can manifest as poor inter- and intra-observer concordance, and this has been the subject of numerous studies [4-8]. While concordance in visual quantitative scoring can be improved by the development of standardized scoring guidelines and extensive training [9, 10], the labor-intensive aspect and the limited dynamic range still remain as major impediments to the widespread use of visual quantitative scoring of IHC images.

Digital image analysis (DIA) based tools overcome some of these limitations of visual quantitative scoring by enabling fast, objective, and highly reproducible quantification of biomarkers from whole-slide IHC images [1, 11]. DIA endpoints are typically continuous variables (e.g. cell density and % positive cells) and offer adequate dynamic range to represent biomarker expression in the IHC image. One of the widely used endpoints to quantify biomarker expression is the H-score [2, 12]. In the H-score algorithm (Figure 1A) individual cells and their sub-cellular compartments (i.e. nucleus, cytoplasm, and cell membrane) are first detected, and based on the relative expression of the biomarker of interest in one or more sub-cellular compartments the cells are classified as either positive or negative. The positive cells are further classified into high (3+), medium (2+), or low (1+) based on the biomarker signal intensity. The H-score is given by the ratio of the weighted sum of the number of positive cells to the total number of detected cells. The H-score captures both the intensity and the proportion of the biomarker of interest from the IHC image and comprises values between 0 and 300, thereby offering a dynamic range to quantify biomarker abundance. A different scoring method developed to quantify estrogen and progesterone receptors in breast cancers, the Allred score [2, 12], assigns separate categorical scores for the intensity (0-3) and the proportion (0-5) of the biomarkers in immunolabeled specimens, and the final score is the sum of these two scores. Compared to the H-score, the Allred score has a limited dynamic range (0-8) and is not extensively used for purposes other than ER/PR quantification in breast cancer. From a digital image analysis standpoint, both the H-score and the Allred score require the detection of individual cells, and this requires robust nucleus and cell segmentation algorithms for individual nucleus detection and delineation of individual cell boundaries.

**Figure 1:**
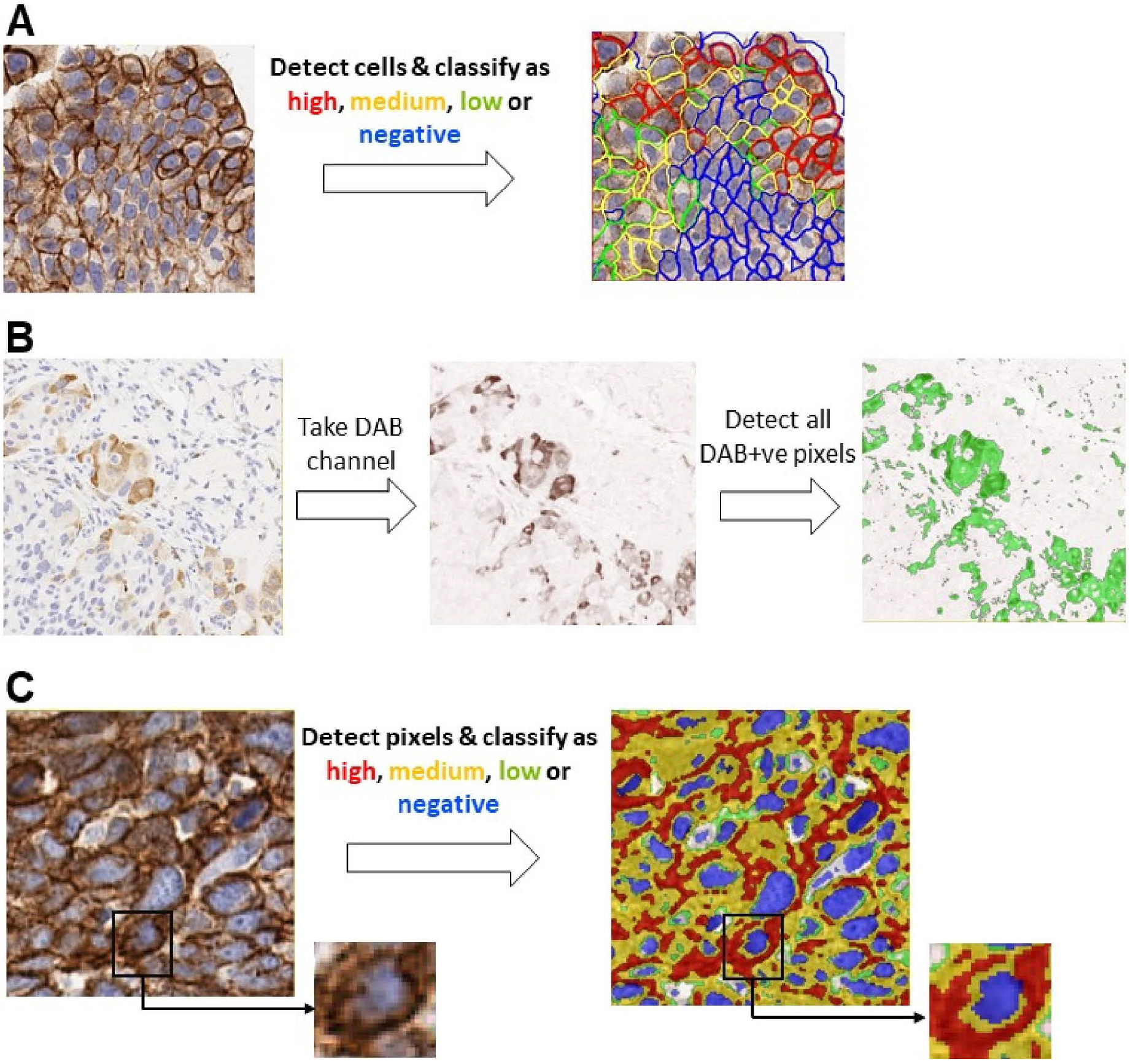
Overview of the different scoring algorithms. Panel A shows the traditional cell-based H-score, panel B shows the average threshold method (ATM) score, and panel C shows the pix H-score.

Another scoring methodology, the average threshold method (ATM), adopts a pixelwise approach for quantifying biomarker abundance [13]. The ATM score does not require the detection of individual nuclei or cells and is solely based on the pixel intensities of the DAB chromogen in the spectrally deconvolved image. Consequently, the calculation of the ATM score is relatively straightforward but at the expense of decreased dynamic range as compared to the H-score.

The AQUA score [14] also makes use of a pixelwise strategy for quantifying biomarker expression. Here, the tissue is fluorescently labeled for the biomarker of interest along with a nuclear stain and a cell membrane marker. This in turn allows the generation of pixel masks pertaining to different subcellular compartments (e.g., cell membrane, nucleus, or cytosolic mask). The AQUA score is then calculated by taking the total fluorescence signal of the biomarker of interest for a given subcellular mask (e.g. the cell-membrane mask) and normalizing it by the total area of the mask [14]. The advantage of the AQUA score is that it offers a broad dynamic range. However, the calculation of the AQUA score requires the development of a fluorescence-based multiplex assay which can be time consuming and technically challenging. Moreover, the use of fluorescence readout masks anatomic and morphological information (e.g. necrotic regions, stroma, etc.) that are readily detectable from a brightfield IHC image.

In this manuscript, three different scoring methods are compared, which are illustrated in Figure 1. We introduce a new DIA method, the pixelwise H-score (pix H-score), for quantifying biomarker abundance from brightfield IHC images by making use of individual pixel intensities in DAB and hematoxylin channels and leveraging weighted intensity averages. Our motivation behind developing the pix H-score is to create a simple, yet robust metric to accurately quantify biomarker expression without relying on the detection and delineation of individual cells and their sub-cellular compartments. The latter makes the implementation of the pix H-score to be relatively straightforward. The pix H-score can be thought of as an equivalent of the traditional H-score that is applied to pixels rather than to cells. The pix H-score takes values between 0 and 300 thereby providing a dynamic range similar to that of the H-score.

We evaluated the performance of pix H-score using IHC images of three different membrane biomarkers P-cadherin, PD-L1, and 5T4. For comparison, we also calculated the ATM score and the DIA H-score for these images, where the latter is a DIA implementation of the traditional H-score. Using the pathologist H-score and biomarker mRNA transcript level (measured using qRT-PCR or NanoString analysis of mRNA in adjacent serial sections) as orthogonal measurements of biomarker abundance, we demonstrate that the pix H-score is either comparable or superior to other DIA endpoints in quantifying biomarker abundance in IHC images. We present the detailed implementation of the pix H-score in two commercial, whole-slide, image analysis software packages, Visiopharm and HALO. We also present an empirical resampling approach to quantitatively assess the ability of the pix H-score to estimate biomarker abundance when it is calculated from select regions within the tumor resection when compared to the whole slide pix H-score. We anticipate that the new metric will have broad applicability and pave the way towards establishing an objective, reproducible strategy to quantify biomarker abundance in IHC images.

## MATERIALS AND METHODS

Previously-developed IHC assays for P-cadherin, PD-L1, and 5T4 were used to immunolabel three cohorts of human tumors. Serial sections from these cohorts were also evaluated for target mRNA via NanoString (P-cadherin and PD-L1) or qRT-PCR (5T4). Following H-scoring of the immunolabeled tumor sections by a pathologist, the concordance between the H-score and mRNA values was evaluated by Spearman correlation. To automate the scoring process through digital image analysis, we implemented several DIA strategies using different software tools. Specifically, we implemented digital H-scoring using QuPath and HALO software packages, the ATM score using Visiopharm software, and the pix H-score, the new digital scoring method, using HALO and Visopharm software packages. To assess the performance of the various DIA algorithms, we calculated the Spearman’s correlation coefficient between each DIA endpoint and two different measurements of biomarker abundance, i.e. the pathologist H-score and the target transcript level as assessed using either NanoString technology or qRT-PCR.

### Immunohistochemistry

All tumor samples used in this study were anonymized specimens from commercial and academic sources that collected the specimens with donor consent under Institutional Review Board-approved procedures. For PD-L1, we used twenty-four cases of routinely collected non-small cell lung carcinoma surgical resections. The SP142 clone of anti-PD-L1 antibody was used as per the manufacturer-recommended protocol. For P-cadherin, we used thirty cases of routinely collected head and neck tumor resections. The P-cadherin IHC assay was developed and optimized on the Dako Autostainer system using a custom anti-P-cadherin antibody that was generated as an analyte specific reagent for use in a clinical diagnostic assay. For 5T4, we used twenty-one cases of routinely collected non-small cell lung tumor resections. The development and validation of the 5T4 IHC assay was reported previously [15]. In all three IHC assays hematoxylin was used as the nuclear counterstain and diaminobenzidine (DAB) was the chromogen that was used to detect the biomarker of interest. P-cadherin and PD-L1 slides were scanned using a Leica Aperio AT2 whole-slide scanner at 20x magnification, whereas 5T4 slides were scanned using a Hamamatsu Nanozoomer whole-slide scanner at 20x magnification.

### NanoString assay

Messenger RNA (mRNA) was isolated from two 4-micron FFPE slide sections using FormaPure® nucleic acid isolation kit according to manufacturer’s instructions with the addition of a DNA digestion step. NanoString technology was used to measure RNA transcript levels using the nCounter assay according to manufacturer’s recommended protocols. Custom nCounter CodeSet containing either the CDH3 probe (for P-cadherin) or the CD274 probe (for PD-L1) was used. One hundred nanograms of total RNA was hybridized to the custom panel for 16 to 20 hours at 65°C. Samples were processed using an automated nCounter sample prep station. Cartridges containing immobilized and aligned reporter complex were subsequently imaged and counted on an nCounter Digital Analyzer set for maximum fields of view. Reporter counts were analyzed and normalized using NanoString nSolver Analysis Software. Briefly, raw counts were multiplied by scaling factors proportional to the sum of counts for spiked in positive control probes to account for individual assay efficiency variation, and to the geometric average of the housekeeping gene probes to account for variability in the mRNA content. FFPE sample sets were normalized to the following housekeeping genes; for P-cadherin: FTL, GAPDH, GUSB, HMBS, HPRT1, OAZ1, PCBP1, PFN1, PPIA, PSAP and TBP; and for PD-L1: AMMECR1L, CNOT10, CNOT4, COG7, DDX50, EDC3, EIF2B4, ERCC3, FCF1, FTL, GPATCH3, GUSB, HDAC3, HPRT1, MTMR14, PPIA, SAP130, TBP, TMUB2, and ZNF143.

### qRT-PCR assay

The qRT-PCR reaction was performed using the TaqMan Probe-Based Gene Expression Analysis and ABI ViiA7 Real-Time PCR Systems (Life Technologies) as described previously [15]. Target gene and endogenous controls were run in quadruplicate for each probe set on prefabricated TaqMan low density array cards. For each tumor sample 1000 ng of cDNA was diluted to 55 uL with nuclease-free water and 55 uL of TaqMan gene expression master mix was added (Life Technologies, cat # 4352042). A total of 100 uL of sample was added to each of the 8 ports on a single card, after which the plate was sealed and centrifuged two times in Sorvall/Heraeus buckets based on manufacturer’s directions. TaqMan array cards were then sealed and loaded into the ABI ViiA7 thermal cycler and run. Default thermal cycling conditions were as follows; the RT-PCR reaction was run on the thermal cycler in three stages; 2 minutes at 50°C, 10 minutes at 90°C and 40 cycles of 15 seconds at 90°C followed by 1 minute at 60°C. ExpressionSuite Software v1.0.3 (Life Technologies) was used to generate automated threshold values for signal amplification for a majority of samples. Rarely were automated thresholds adjusted manually. Amplification plots resulting in Ct values >35 were discarded, as were those plots that generated a Ct value but did not display a trend of logarithmic amplification. All Ct values were exported from the ExpressionSuite software and relative quantification calculations were performed in Microsoft Excel 2010.

### Digital Image analysis

IHC images of P-cadherin, PD-L1, and 5T4 were analyzed at 20x magnification using multiple software packages. The detailed implementation in each software package is described below. Briefly, the traditional cell-based H-score was implemented in HALO (Version 2.3) and QuPath (Version 0.2.0-m2) and was calculated based on the cell-membrane localized biomarker signal. The ATM score was implemented in Visiopharm (Version 2017.7.3.4069) and the pix H-score was implemented in Visiopharm and HALO.

### HALO implementation of H-score (H-score (HALO))

The membrane algorithm (v1.4) in HALO was used to detect cells and calculate the H-score. The algorithm first deconvolves the IHC image into hematoxylin and DAB channels, then detects individual cells and their subcellular compartments, i.e. nucleus and cell membrane, in the image, and scores the cells as high, medium, and low based on the average DAB signal associated with the cell membrane. The thresholds for high, medium, and low were determined separately for each biomarker by examining the membrane-associated DAB signal across multiple images pertaining to that biomarker. A separate algorithm was implemented for each biomarker in order to optimize the detection and segmentation of the nucleus and cell membrane specific to that biomarker. The App outputs the number of negative, high, medium and low cells, which is then used to calculate the H-score that is given by

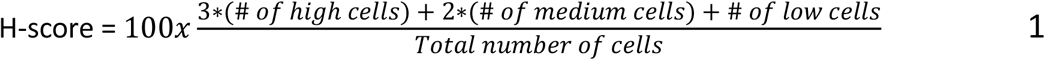

### QuPath implementation of H-score (H-score (QuPath))

QuPath is an open-source software for whole-slide image analysis of histopathology data [16]. A script was written in the Groovy programming language to detect cells and score them as high, medium, and low based on the average DAB signal in the cell membrane. The script first deconvolves the IHC image into hematoxylin and DAB channels. A watershed-based cell and membrane detection algorithm (Analyze -> Cell Analysis -> Cell + membrane detection) was used to detect individual cells and identify their subcellular compartments, i.e. nucleus and cell membrane. The cell detection algorithm includes a pre-processing step that involves a local background subtraction by using the minimum filter. The optional median filtering step was not used. Cells that were devoid of a nucleus (due to weak or missing hematoxylin staining) were excluded and the remaining cells were scored as high, medium, and low based on the mean DAB signal associated with the membrane compartment. The thresholds for high, medium, and low were determined separately for each biomarker. A separate script was implemented for each biomarker in order to optimize the detection and segmentation of the nucleus and cell membrane specific to that biomarker. The script outputs the total number detected cells along with the number of high, medium, and low cells, which is then used to calculate the H-score that is given in Eq. 1.

### ATM score

The motivation behind the ATM score is discussed elsewhere [13]. Briefly, the idea is to use all the intensity values in the DAB channel so that the final metric is independent of the choice of the thresholds. Further, the ATM score is a pixel-based metric that does not depend on the detection of individual cells and/or its subcellular components. Assuming 8-bit resolution for the color-deconvolved biomarker channel, the ATM score is given by [13]

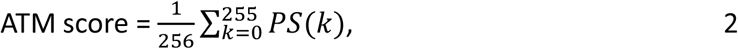

where PS(k) denotes the proportion of pixels with intensity greater than or equal to k, where k takes values from 0 to 255 (i.e. 2^8^ grey levels). If n denotes the total number of pixels in the biomarker channel, b_i_ denotes the biomarker intensity at the i^th^ pixel for I = 1,..,n, and I(b_i_ > k) denotes an indicator function, i.e. I(b_i_ > k) = 1 if b_i_ > k and 0 otherwise, then the term PS(k) can be written as

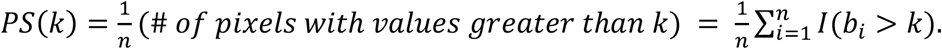

Substituting the above equation in Eq. 2, we have

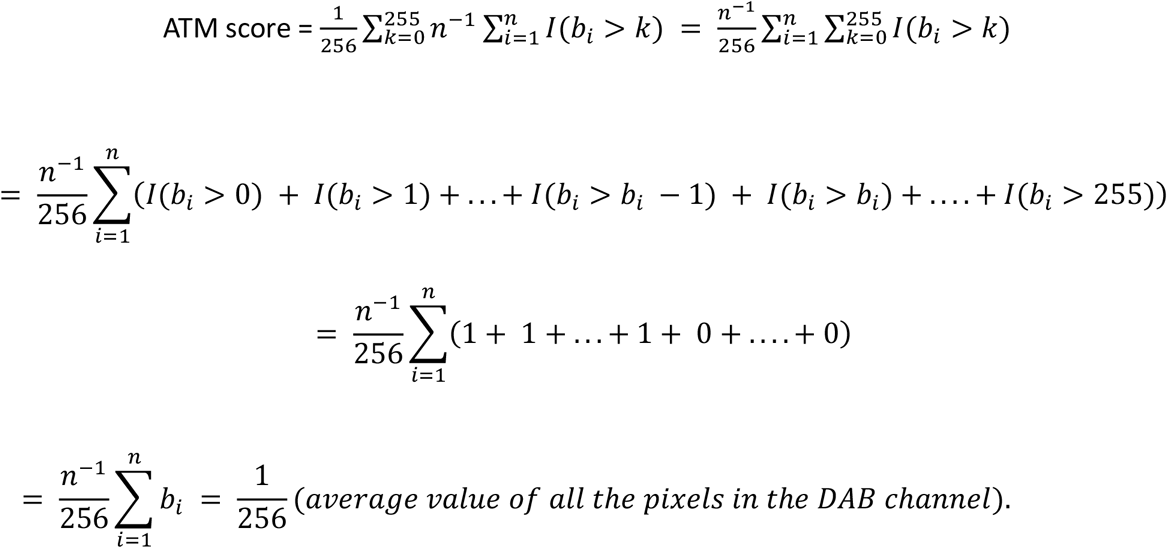

From the above equation, we see that the ATM score is a weighted average of all the pixels in the DAB channel. The ATM score was implemented in Visiopharm software. The IHC image was color deconvolved into hematoxylin and DAB channels. Therefore, the ATM score was calculated by taking the average intensity of all DAB positive pixels and then dividing this by 256.

### Visiopharm implementation of pix H-score (pix H-score (VIS))

A threshold-based detection App was used to implement the pix H-score in Visiopharm. The App first deconvolves the IHC image into hematoxylin and DAB channels. The App then detects and classifies DAB positive pixels as high, medium, and low, and then detects the hematoxylin positive pixels. The thresholds for DAB and hematoxylin were separately selected for each biomarker. The App then outputs the total area of the DAB high, DAB medium, and DAB low pixels and the hematoxylin positive pixels. These values are then used to calculate the pix H-score which is given by:

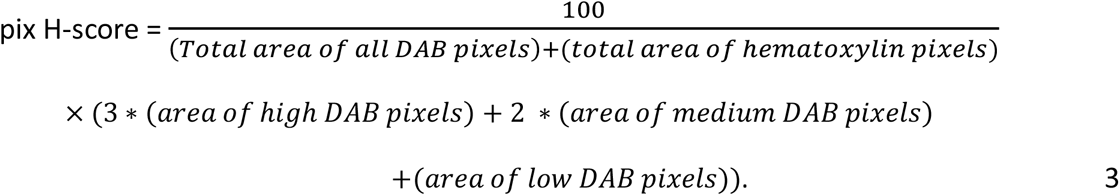

In Visiopharm, the intensity-based thresholding algorithm depends on the order in which the different color-deconvolved channels are specified. For instance, if a pixel contains both hematoxylin and DAB signal that are above their respective threshold values for positivity and the DAB channel is first analyzed followed by the hematoxylin channel, then that pixel will be labeled as positive only for the DAB channel. In other words, if a pixel is found to be positive for one of the color-deconvolved channels then it is excluded from any subsequent classification for the other color-deconvolved channels.

### HALO implementation of pix H-score (pix H-score (HALO))

The area quantification algorithm (v2.1.3) in HALO was used to calculate the pix H-score with the number of phenotypes set to 1. The algorithm deconvolves the IHC image into hematoxylin and DAB channels and can detect and classify hematoxylin and DAB positive pixels as high, medium, and low based on a user defined threshold. For the calculation of pix H-score, a single threshold was used to detect all hematoxylin positive pixels and three separate thresholds were used to detect and classify the DAB positive pixels. In HALO, these thresholds take values between 0 and 1. In order to keep the thresholds implemented in Visiopharm and HALO identical, the threshold values used in Visiopharm, which take values between 0 – 255, were rescaled to take values between 0 and 1 and these were then used in HALO. Unlike Visiopharm, HALO keeps track of the detected pixels in the DAB and hematoxylin channels separately. Consequently, pixels that contain both DAB and hematoxylin signal that are above the thresholds will be accounted for in both the hematoxylin and DAB channels. In order to mimic the Visiopharm implementation of pix H-score, we define a third channel, which is denoted as phenotype 1 channel in HALO that pertains to pixels that are positive for hematoxylin but negative for DAB. This phenotype 1 channel will contain pixels that are analogous to the hematoxylin positive pixels detected in the Visiopharm implementation of pix H-score algorithm. The algorithm outputs the area high, medium, and low pixels in the DAB channel, the area of positive pixels in the phenotype 1 channels, which is used to as an estimate of the total area of pixels containing only the hematoxylin signal. These values are then used in Eq. 3 to calculate the pix H-score.

### Statistical analysis

Spearman’s rank correlation coefficient was calculated to assess the correlation between the DIA endpoint and biomarker abundance. The William’s t test was used to test for significant difference between a pair of dependent correlation coefficients [17, 18].

### Spatial resampling analysis

For each biomarker, an empirical resampling procedure was performed on every whole-slide IHC image. The viable tissue region was sampled by non-overlapping circular regions of radius 0.8 mm (Figure 6A). For each region, the area of DAB high, DAB medium, DAB low, and hematoxylin positive pixels were determined using Visiopharm. The results were exported to MATLAB (Mathworks, Natick, MA) for subsequent analysis. For every IHC image, N different circular regions were randomly selected (N = 1 – 50), and a regional pix H-score was calculated using the area of DAB high pixels, DAB medium pixels, DAB low pixels, and hematoxylin positive pixels that were summed from the N circular regions. This procedure is repeated N_iter_ times with replacement (N_iter_ = 100 for all the biomarkers). Then for each iteration k = 1,…,N_iter_, the Spearman correlation coefficient C(N,k) is computed between the regional pix H-score and the corresponding pathologist H-score. The average Spearman correlation coefficient for each value of N is computed using the formula

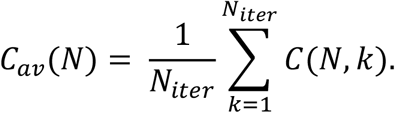

## RESULTS

### DIA algorithms for P-cadherin quantification

IHC images for P-cadherin (Figure 2A) showed strong immunoreactivity at the cell membrane and in the cytoplasm, which was consistent with prior reports [19, 20]. Spearman’s correlation analysis of the membrane H-scores of the 30 cases immunolabeled for P-cadherin, as assessed by a board-certified pathologist (see Supplementary Table 1), and NanoString nCounter values for P-cadherin mRNA transcript from serial sections of the same cases had a correlation coefficient of 0.81, p<0.0001 (Figure 2B).

**Figure 2.**
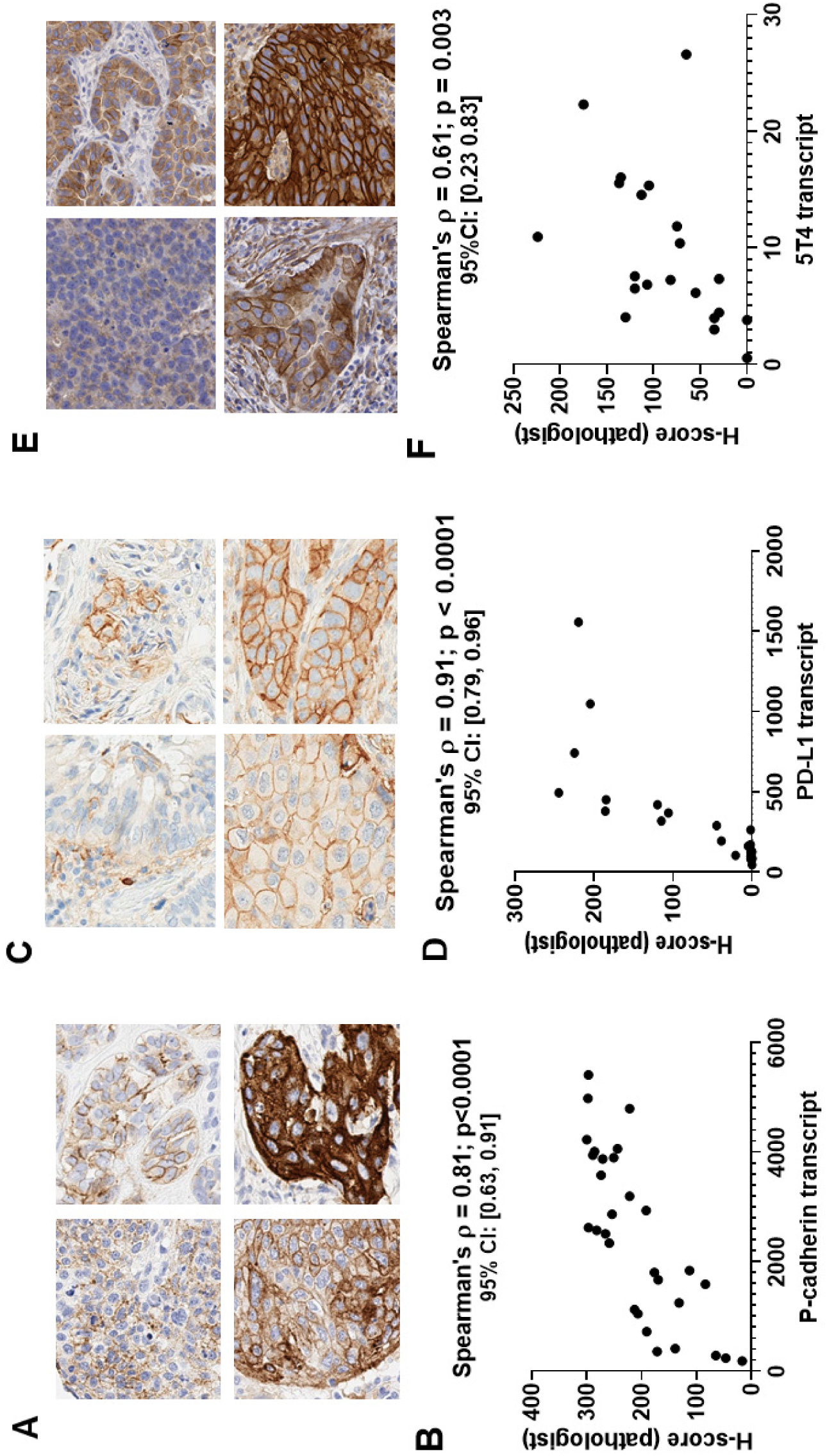
P-cadherin, PD-L1 and 5T4 IHC datasets. Panels A, C and E show representative images at 20x magnification with varying levels of P-cadherin, PD-L1 and 5T4 expression, respectively, in tumor resections. Panels B, D and F show the plot of the pathologist H-score versus mRNA transcript level for P-cadherin (n = 30 cases), PD-L1 (n = 24 cases) and 5T4 (n = 21 cases), respectively. The panels also show Spearman’s correlation coefficient along with the p-value and 95% confidence interval.

When compared to the P-cadherin pathologist H-score, all P-cadherin DIA endpoints yielded positive correlations (Figures 3A-3E). The correlation with the ATM score (Figure 3C) and pix H-score (Figures 3D and 3E) were higher than the correlations with the DIA based H-scores (Figures 3A and 3B). More specifically, the Spearman’s correlation coefficient for HALO and QuPath DIA H-scores were 0.5 (p = 0.005) and 0.39 (p = 0.03), respectively, whereas the Spearman’s correlation coefficient for the ATM score, the VIS pix H-score and the HALO pix H-score were 0.78 (p<0.001), 0.77 (p<0.0001) and 0.88 (p<0.0001), respectively. When compared to the P-cadherin transcript, all DIA endpoints similarly yielded positive correlations (Figures 3F-3J), with the pix H-score exhibiting the highest Spearman’s correlation coefficient (Figures 3I and 3J; ρ = 0.83 and ρ = 0.81, respectively, for VIS and HALO pix H-score; p < 0.0001) followed by the ATM score (Figure 3H; ρ = 0.62, p < 0.0001) and the DIA H-scores (Figures 3F and 3G; ρ = 0.5, p = 0.005 for HALO and ρ = 0.45, p = 0.01 for QuPath).

**Figure 3.**
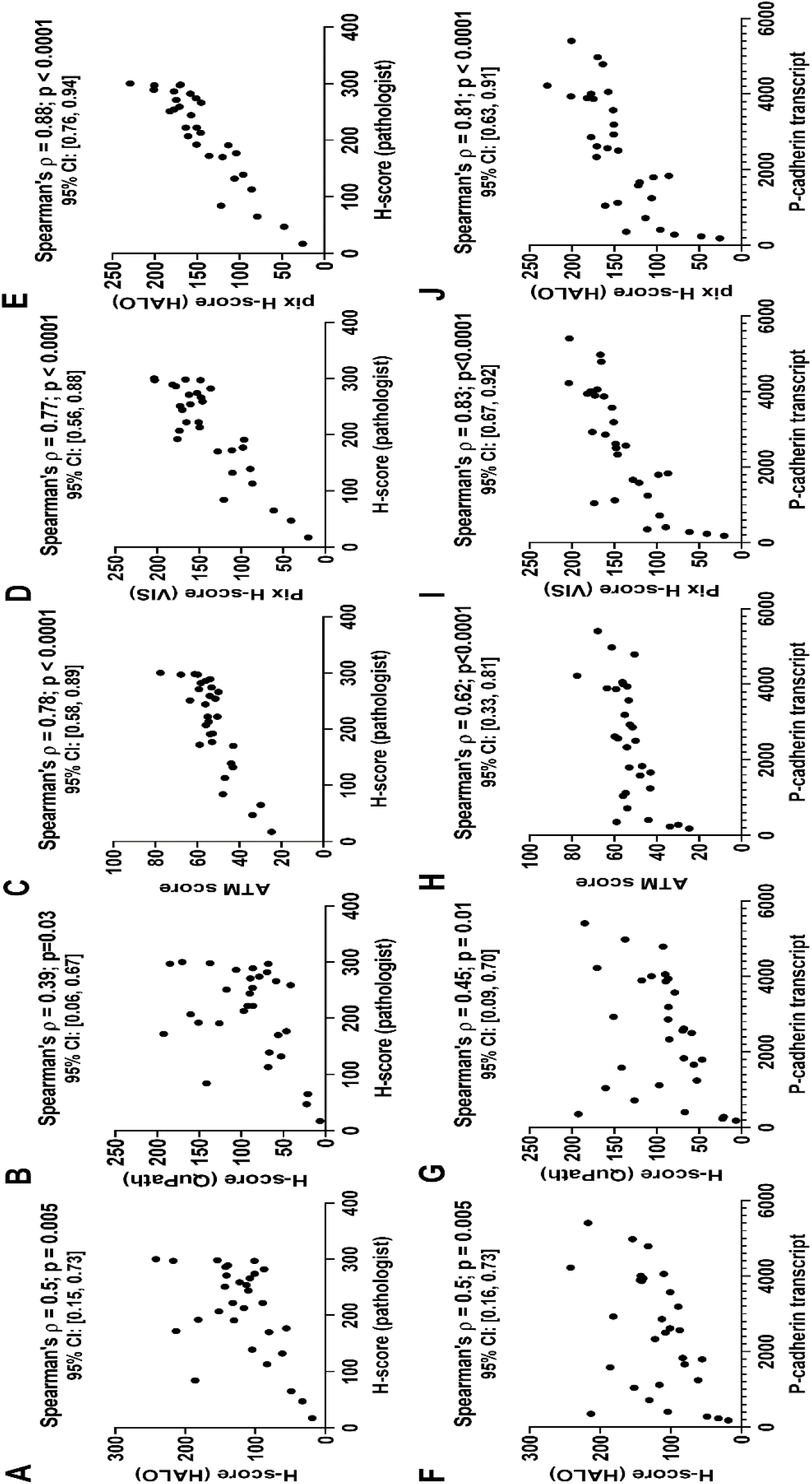
Performance of DIA endpoints obtained from P-cadherin IHC images. Panels A through E show the plots of different DIA endpoints versus pathologist H-score for a cohort of 30 head and neck cancer resections. Panels F through J show the plots of different DIA endpoints versus P-cadherin mRNA transcript for the same 30 cases. Each panel also shows the Spearman’s correlation coefficient between the two quantities plotted in that panel along with the p-value and the 95% confidence interval.

We next investigated whether the differences in the Spearman correlation coefficients for the various DIA endpoints are statistically significant. Table 1 shows the results of our statistical analysis where we carried out pairwise comparisons of the correlation coefficients for different DIA endpoints obtained from P-cadherin IHC images. Our analysis shows that the correlation coefficient between the pix H-score and either of the biomarker abundance endpoints (pathologist H-score and P-cadherin transcript) is significantly higher than the correlation coefficient between DIA based H-scores and biomarker abundance endpoints. This suggests that for the P-cadherin dataset, the pix H-score is a better DIA metric to quantify biomarker abundance over traditional DIA based H-score. In the case of the ATM score, we observe a mixed result in that the correlation coefficient between pix H-score and P-cadherin transcript is significantly higher than the correlation coefficient between ATM score and P-cadherin transcript, whereas statistical significance is lost when we consider the pathologist H-score as the reference for biomarker abundance (Table 1). We also compared the two DIA based H-scores. We found no significant difference in the Spearman’s correlation coefficient between QuPath H-score and biomarker abundance endpoints versus HALO H-score and biomarker abundance endpoints (Table 1). Similarly, we found no significant difference in the correlation coefficients for the HALO and VIS implementations of the pix H-score for P-cadherin.

**Table 1.** Table lists the results of William’s t-test to test for significant difference in the Spearman correlation coefficients between P-cadherin transcript or pathologist H-score and different DIA endpoints.

| Statistical analysis of correlation coefficients for P-cadherin | $\rho_{12}$ | $\rho_{13}$ | z-score | p-value | Result |
| --- | --- | --- | --- | --- | --- |
| Comparing pairwise correlations between DIA endpoint and Pathologist H-score |  |  |  |  |  |
| $\rho(\text{Path H-score, H-score HALO})$ vs $\rho(\text{Path H-score, H-score QuPath})$ | 0.50 | 0.39 | 1.48 | 0.15 | No significant difference |
| $\rho(\text{Path H-score, pix H-score VIS})$ vs $\rho(\text{Path Hscore, pix H-score HALO})$ | 0.77 | 0.88 | -2.5 | 0.01 | Significant difference |
| $\rho(\text{Path H-score, pix H-score VIS})$ vs $\rho(\text{Path H-score, H-score HALO})$ | 0.77 | 0.50 | 3.01 | 0.005 | Significant difference |
| $\rho(\text{Path H-score, pix H-score VIS})$ vs $\rho(\text{Path H-score, H-score QuPath})$ | 0.77 | 0.39 | 3.87 | 0.0006 | Significant difference |
| $\rho(\text{Path H-score, pix H-score VIS})$ vs $\rho(\text{Path H-score, ATM score})$ | 0.77 | 0.78 | -0.12 | 0.90 | No significant difference |
| $\rho(\text{Path H-score, pix H-score HALO})$ vs $\rho(\text{Path H-score, H-score HALO})$ | 0.88 | 0.50 | 5.21 | 1.7e-05 | Significant difference |
| $\rho(\text{Path H-score, pix H-score HALO})$ vs $\rho(\text{Path H-score, H-score QuPath})$ | 0.88 | 0.39 | 5.34 | 1.2E-05 | Significant difference |
| $\rho(\text{Path H-score, pix H-score HALO})$ vs $\rho(\text{Path H-score, ATM score})$ | 0.88 | 0.78 | 1.64 | 0.11 | No significant difference |
| Comparing pairwise correlations between DIA endpoint and P-cadherin transcript |  |  |  |  |  |
| $\rho(\text{Pcad transcript, H-score HALO})$ vs $\rho(\text{Pcad transcript, H-score QuPath})$ | 0.50 | 0.45 | 0.79 | 0.43 | No significant difference |
| $\rho(\text{Pcad transcript, pix H-score VIS})$ vs $\rho(\text{Pcad transcript, pix H-score HALO})$ | 0.83 | 0.81 | 0.50 | 0.62 | No significant difference |
| $\rho(\text{Pcad transcript, pix H-score VIS})$ vs $\rho(\text{Pcad transcript, H-score HALO})$ | 0.83 | 0.5 | 4.19 | 0.0002 | Significant difference |
| $\rho(\text{Pcad transcript, pix H-score VIS})$ vs $\rho(\text{Pcad transcript, H-score QuPath})$ | 0.83 | 0.45 | 4.49 | 0.0001 | Significant difference |
| $\rho(\text{Pcad transcript, pix H-score VIS})$ vs $\rho(\text{Pcad transcript, ATM score})$ | 0.83 | 0.62 | 2.45 | 0.02 | Significant difference |
| $\rho(\text{Pcad transcript, pix H-score HALO})$ vs $\rho(\text{Pcad transcript, H-score HALO})$ | 0.81 | 0.5 | 3.31 | 0.002 | Significant difference |
| $\rho(\text{Pcad transcript, pix H-score HALO})$ vs $\rho(\text{Pcad transcript, H-score QuPath})$ | 0.81 | 0.45 | 3.22 | 0.003 | Significant difference |
| $\rho(\text{Pcad transcript, pix H-score HALO})$ vs $\rho(\text{Pcad transcript, ATM score})$ | 0.81 | 0.62 | 2.37 | 0.025 | Significant difference |

### DIA algorithms for PD-L1 quantification

IHC images for PD-L1 (Figure 2C) showed strong immunoreactivity at the cell membrane and minimal to no cytoplasmic staining, which was consistent with prior reports [19, 20]. Spearman’s correlation analysis of the membrane H-scores of the 24 cases immunolabeled for PD-L1, as assessed by a board-certified pathologist (see Supplementary Table 1), and NanoString nCounter values for PD-L1 mRNA transcript from serial sections of the same cases had a correlation coefficient of 0.91, p<0.0001 (Figure 2D).

When compared to the pathologist H-score, all DIA endpoints yielded positive correlations (Figures 4A-4E). The Spearman’s correlation coefficient for the HALO H-score, QuPath H-score, ATM score, VIS pix H-score and HALO pix H-score with respect to the pathologist H-score were 0.69 (p=0.0002), 0.74 (p<0.0001), 0.55 (p = 0.005), 0.76 (p < 0.0001) and 0.71 (p < 0.0001), respectively. When compared to the PD-L1 transcript, all DIA endpoints similarly yielded positive correlations (Figures 4F-4J). The Spearman’s correlation coefficient for the HALO H-score, QuPath H-score, ATM score, VIS pix H-score and HALO pix H-score with respect to PD-L1 transcript were 0.73 (p<0.0001), 0.75 (p<0.0001), 0.55 (p = 0.005), 0.79 (p<0.0001) and 0.79 (p<0.0001), respectively.

**Figure 4.**
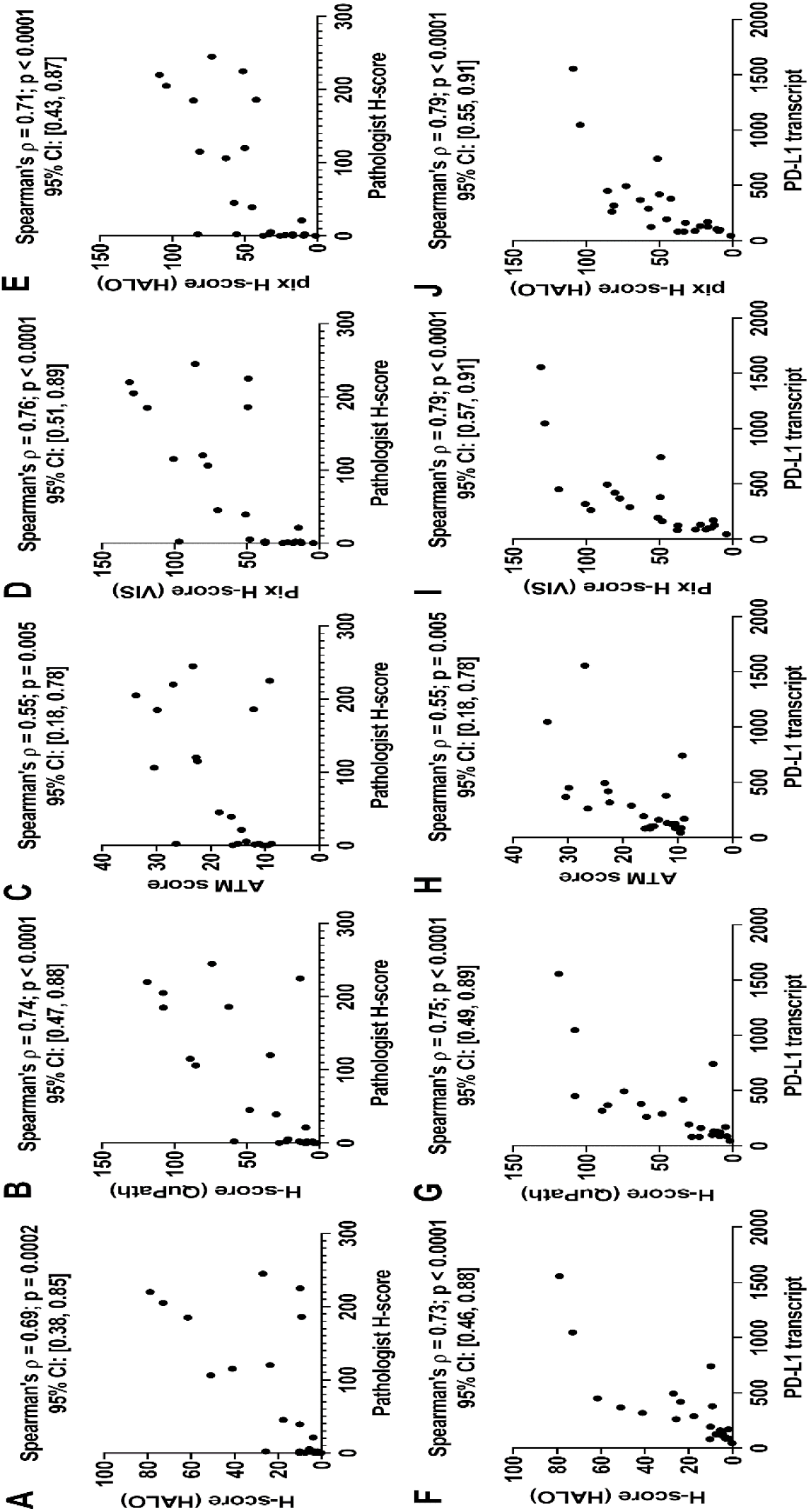
Performance of DIA endpoints obtained from PD-L1 IHC images. Panels A-E show plots of the different DIA endpoints as a function of the pathologist H-score, while panels F-J show the same as a function of PD-L1 mRNA transcript for a cohort of 24 lung cancer resections. All panels show the Spearman’s correlation coefficient between the two quantities plotted in that panel along with the p-value and the 95% confidence interval.

Statistical analysis of the Spearman’s correlation coefficients revealed that there is no significant difference in the correlation coefficient between DIA based H-scores and PD-L1 biomarker abundance endpoints versus the correlation coefficient between pix H-score and PD-L1 biomarker abundance endpoints (Table 2). This shows that the performance of pix H-score is analogous to that of the DIA based H-score which is in contrast with our observations for P-cadherin. Also, there was no significant difference in Spearman’s correlation coefficient between HALO and QuPath implementations of the H-score, which is analogous to what we observed for P-cadherin. In addition, we observed that there was no significant difference between the HALO and Visiopharm implementations of the pix H-score for PD-L1. Spearman’s correlation coefficients between the pix H-score and PD-L1 biomarker abundance endpoints were mostly significantly higher than Spearman’s correlation coefficients between ATM score and PD-L1 biomarker abundance endpoints (Table 2). Although both the pix H-score and the ATM score are pixel-based algorithms, the higher Spearman’s correlation coefficient for the pix H-score suggests that this algorithm is superior to the ATM score in estimating biomarker abundance for PD-L1.

**Table 2.**
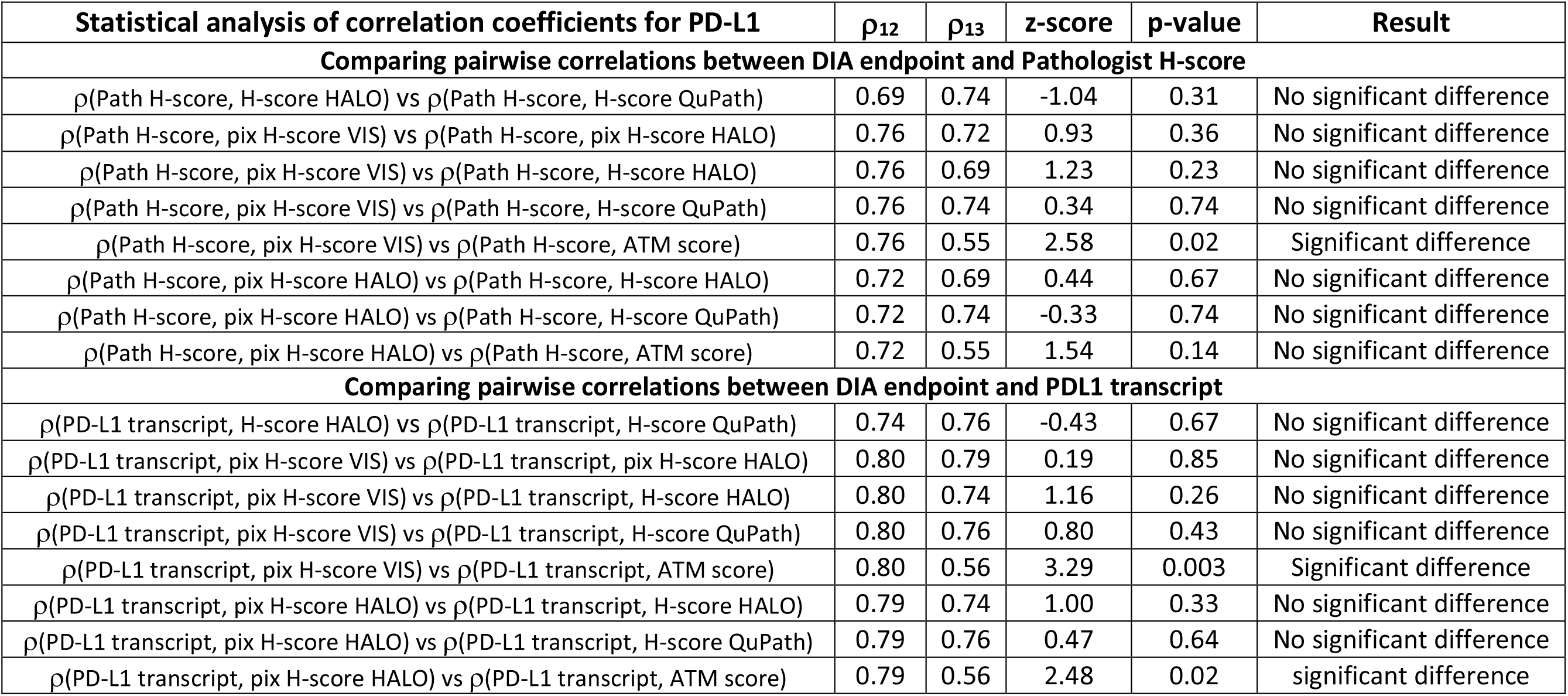
Table lists the results of William’s t-test to test for significant difference in the Spearman correlation coefficients between PD-L1 mRNA transcript or pathologist H-score and different DIA endpoints.

### DIA algorithms for 5T4 quantification

IHC images for 5T4 (Figure 2E) showed strong immunoreactivity at the cell membrane with limited cytoplasmic staining, which was consistent with prior reports [15]. Spearman’s correlation of the membrane H-scores of the 21 cases immunolabeled for 5T4, as assessed by a board-certified pathologist (see Supplementary Table 1), and qRT-PCR values for 5T4 mRNA transcript from serial sections of the same cases had a ρ value of 0.61, p=0.003 (Figure 2F).

When compared to the pathologist H-score, all DIA endpoints yielded positive correlations (Figures 5A-5E). The Spearman’s correlation coefficient for the HALO H-score, QuPath H-score, ATM score, VIS pix H-score and HALO pix H-score with respect to the pathologist H-score were 0.75 (p<0.0001), 0.79 (p<0.0001), 0.76 (p < 0.0001), 0.83 (p < 0.0001) and 0.82 (p < 0.0001), respectively. When compared to the 5T4 transcript, all DIA endpoints similarly yielded positive correlations (Figures 5F-5J). The Spearman’s correlation coefficient for the HALO H-score, Qupath H-score, ATM score, VIS pix H-score and HALO pix H-score with respect to 5T4 transcript were 0.74 (p<0.0001), 0.55 (p=0.01), 0.69 (p = 0.0007), 0.76 (p<0.0001) and 0.74 (p=0.0001), respectively.

**Figure 5.**
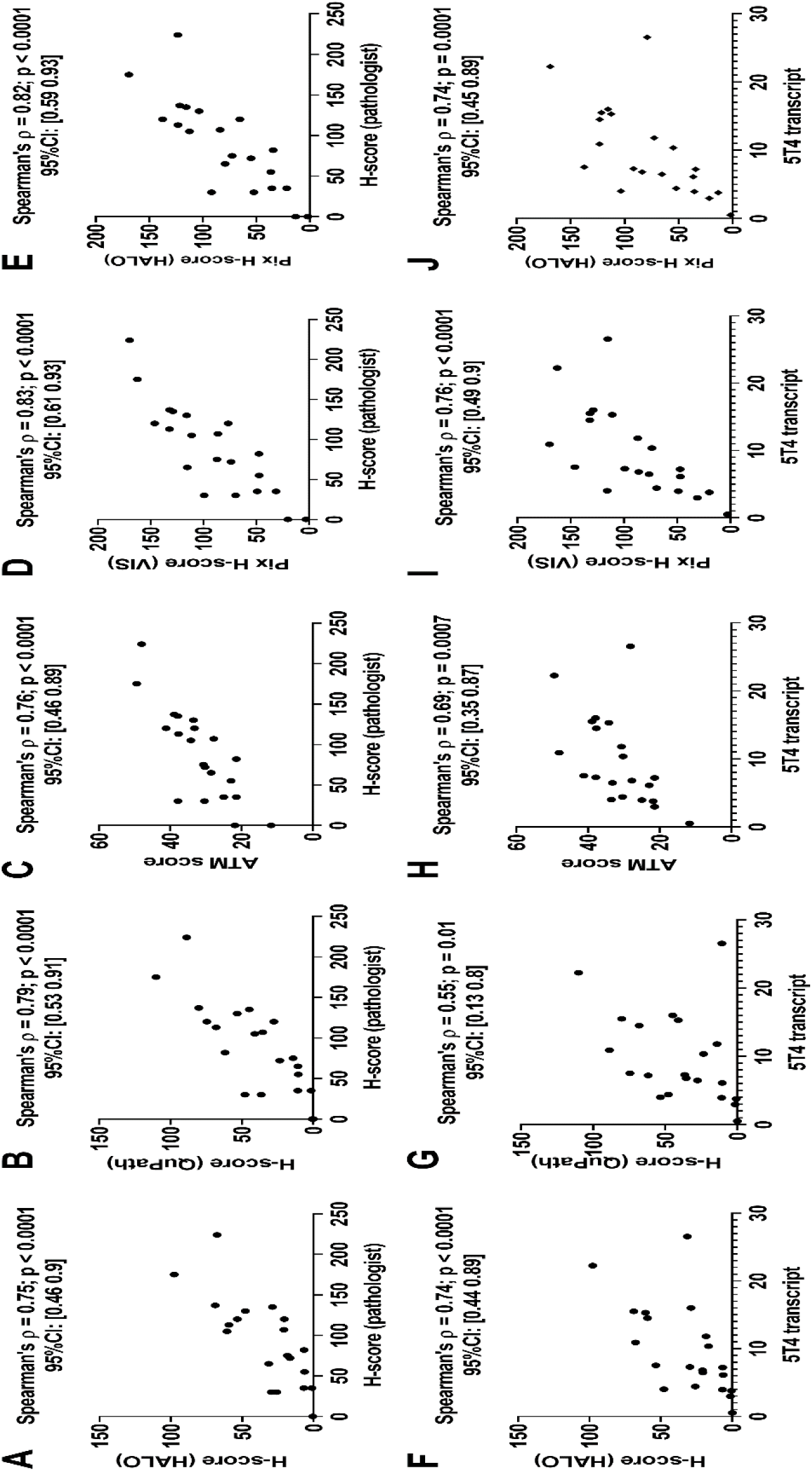
Performance of DIA endpoints obtained from 5T4 IHC images. Panels A-E show plots of the different DIA endpoints as a function of the pathologist H-score, while panels F-J show the same as a function of 5T4 mRNA transcript for a cohort of 21 lung cancer resections. All panels show the Spearman’s correlation coefficient between the two quantities plotted in that panel along with the p-value and the 95% confidence interval.

**Figure 6.**
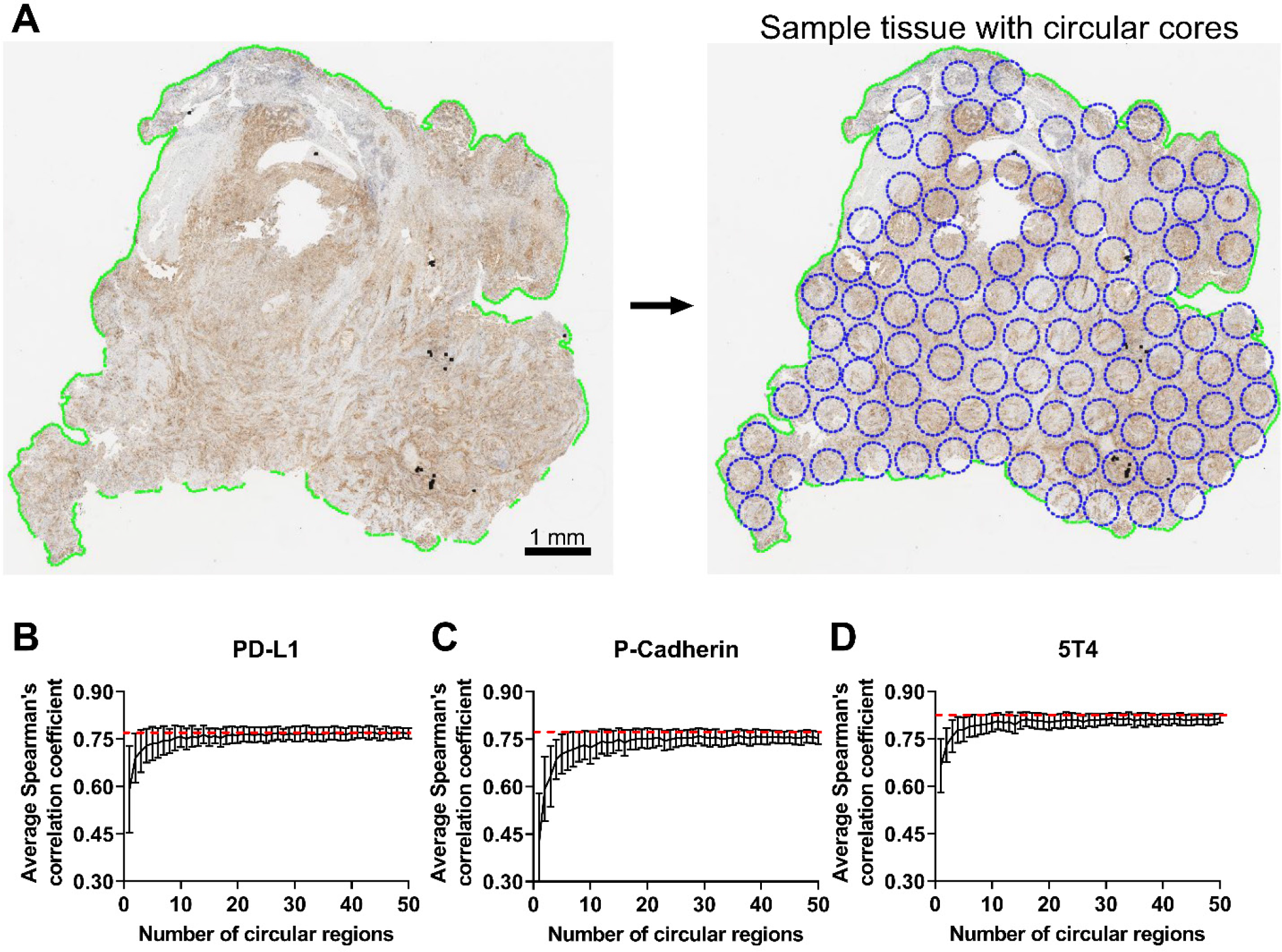
Empirical approach to assess robustness of pix H-score to spatial sampling. Panel A shows the breakup of the tumor resection into non overlapping circular regions. Panels B, C and D show the results of the bootstrap analysis for PD-L1, P-cadherin and 5T4, respectively, where the average Spearman’s Correlation coefficient between the regional pix H-score estimate from N circular regions and pathologist H-score is plotted as a function of the number of circular regions, where N varies from 1 to 50. The red dashed line shows the Spearman’s correlation coefficient between whole-slide Pix H-score and pathologist H-score for that biomarker. Error bars indicate ± SEM.

Statistical analysis of the Spearman’s correlation coefficients revealed that there is no significant difference in the correlation coefficient between each of the DIA based endpoints and pathologist H-score (Table 3). An analogous behavior was also observed for the correlation coefficient between each of the DIA based endpoints and 5T4 transcript except for the QuPath H-score. Specifically, the correlation between QuPath H-score and 5T4 transcript was significantly lower than the correlation between the HALO H-score or the pix H-score endpoints and 5T4 transcript (Table 3). Finally, we note that there is no significant difference in the correlation coefficient between the HALO and Visiopharm implementations of the pix H-score and either of the biomarker abundance endpoints for 5T4. These results suggest that the pix H-score algorithm has comparable performance to the other DIA algorithms to quantify biomarker abundance for 5T4.

**Table 3.** Table lists the results of William’s t-test to test for significant difference in the Spearman correlation coefficients between 5T4 mRNA transcript or pathologist H-score and different DIA endpoints.

| Statistical analysis of correlation coefficients for 5T4 | | $\rho_{12}$ | $\rho_{13}$ | z-score | p-value | Result |
| --- | --- | --- | --- | --- | --- | --- |
| Comparing pairwise correlations between DIA endpoint and Pathologist H-score |  |  |  |  |  |  |
| $\rho(\text{Path H-score, H-score HALO})$ vs $\rho(\text{Path H-score, H-score QuPath})$ | | 0.75 | 0.79 | -0.49 | 0.63 | No significant difference |
| $\rho(\text{Path H-score, pix H-score VIS})$ vs $\rho(\text{Path H-score, pix H-score HALO})$ | | 0.83 | 0.82 | 0.31 | 0.76 | No significant difference |
| $\rho(\text{Path H-score, pix H-score VIS})$ vs $\rho(\text{Path H-score, H-score HALO})$ | | 0.83 | 0.75 | 1.69 | 0.11 | No significant difference |
| $\rho(\text{Path H-score, pix H-score VIS})$ vs $\rho(\text{Path H-score, H-score QuPath})$ | | 0.83 | 0.79 | 0.49 | 0.63 | No significant difference |
| $\rho(\text{Path H-score, pix H-score VIS})$ vs $\rho(\text{Path H-score, ATM score})$ | | 0.83 | 0.76 | 1.37 | 0.18 | No significant difference |
| $\rho(\text{Path H-score, pix H-score HALO})$ vs $\rho(\text{Path H-score, H-score HALO})$ | | 0.82 | 0.75 | 1.46 | 0.16 | No significant difference |
| $\rho(\text{Path H-score, pix H-score HALO})$ vs $\rho(\text{Path H-score, H-score QuPath})$ | | 0.82 | 0.79 | 0.37 | 0.71 | No significant difference |
| $\rho(\text{Path H-score, pix H-score HALO})$ vs $\rho(\text{Path H-score, ATM score})$ | | 0.82 | 0.76 | 1.32 | 0.20 | No significant difference |
| Comparing pairwise correlations between DIA endpoint and 5T4 transcript |  |  |  |  |  |  |
| $\rho(5T4 \text{ transcript, H-score HALO})$ vs $\rho(5T4 \text{ transcript, H-score QuPath})$ | | 0.74 | 0.55 | 2.11 | 0.05 | Significant difference |
| $\rho(5T4 \text{ transcript, pix H-score VIS})$ vs $\rho(5T4 \text{ transcript, pix H-score HALO})$ | | 0.76 | 0.74 | 0.67 | 0.51 | No significant difference |
| $\rho(5T4 \text{ transcript, pix H-score VIS})$ vs $\rho(5T4 \text{ transcript, H-score HALO})$ | | 0.76 | 0.74 | 0.50 | 0.63 | No significant difference |
| $\rho(5T4 \text{ transcript, pix H-score VIS})$ vs $\rho(5T4 \text{ transcript, H-score QuPath})$ | | 0.76 | 0.55 | 2.40 | 0.03 | Significant difference |
| $\rho(5T4 \text{ transcript, pix H-score VIS})$ vs $\rho(5T4 \text{ transcript, ATM score})$ | | 0.76 | 0.69 | 1.40 | 0.18 | No significant difference |
| $\rho(5T4 \text{ transcript, pix H-score HALO})$ vs $\rho(5T4 \text{ transcript, H-score HALO})$ | | 0.74 | 0.74 | 0.10 | 0.92 | No significant difference |
| $\rho(5T4 \text{ transcript, pix H-score HALO})$ vs $\rho(5T4 \text{ transcript, H-score QuPath})$ | | 0.74 | 0.55 | 2.10 | 0.05 | Significant difference |
| $\rho(5T4 \text{ transcript, pix H-score HALO})$ vs $\rho(5T4 \text{ transcript, ATM score})$ | | 0.74 | 0.69 | 1.11 | 0.28 | No significant difference |

### Effect of spatial sampling on pix H-score

We next investigated the robustness of the pix H-score when it is calculated from select regions within the tissue section as opposed to the entire tumor resection. For this purpose, a statistical sampling procedure known as bootstrapping needs to be performed. However, technical limitations in Visiopharm and HALO software packages precluded us from implementing a formal bootstrapping procedure. Therefore, we resorted to an empirical resampling approach (see Methods for details) wherein for a given biomarker each tumor resection was divided into non-overlapping circular regions (Figure 6A). N different circular regions (N ranging from 1 to 50) were randomly selected, and a regional pix H-score was computed from these circular regions. Then the Spearman’s correlation coefficient between the pathologist H-score and the regional pix H-score was computed for that biomarker. This procedure was repeated 100 times for all the tumor resections pertaining to that biomarker, and the average Spearman correlation coefficient from 100 iterations was then plotted as a function of the number of circular regions N.

Figures 6B, 6C and 6D show the behavior of the average Spearman’s correlation coefficient for PD-L1, P-cadherin and 5T4, respectively, between pathologist H-score and the regional pix H-score as a function of the number of circular regions from which the regional pix H-score was calculated. For all the biomarkers, we see that for fewer than five circular regions the average Spearman correlation coefficient between the regional pix H-score and pathologist H-score is consistently smaller than the Spearman’s correlation coefficient between the whole-slide pix H-score and pathologist H-score (shown by the red dashed line). When 10 or more circular regions are sampled the average Spearman’s correlation coefficient for the regional pix H-score starts to plateau out and reaches a steady state. In the case of PD-L1, the plateau region converges with the Spearman’s correlation coefficient between the whole-slide pix H-score and pathologist H-score (Figure 6B). In contrast, for P-cadherin 5T4 the plateau region is slightly lower than the Spearman’s correlation coefficient for the whole-slide pix H-score (Figures 6C and 6D). A similar behavior is also observed when biomarker mRNA levels are used as the reference ground truth data in the Spearman’s correlation coefficient calculation (data not shown).

## DISCUSSION

Robust quantification of biomarker expression in tissue sections is a critical need in many diagnostic and investigative pathology workflows. Our motivation to develop a new digital image analysis metric was driven by the need to automate the process of manual scoring by a pathologist. Digital image analysis holds the promise to offer a fast, objective, and reproducible strategy to quantify biomarker expression from histopathology images. In this manuscript, we introduced an unsupervised algorithm, the pix H-score. With it we quantified P-cadherin, PD-L1, and 5T4 signals in immunolabeled FFPE sections of human tumors and found good correlation between the digitally-analyzed IHC signals and manual (visual) signal quantitation as performed by a board certified pathologist. As pathologist scoring is known to be susceptible to intra- and inter-observer variability, we also used biomarker mRNA level as an orthogonal measurement of biomarker abundance to validate the pix H-score. Our observation that there was good concordance between both digital and visual IHC signal quantitation and mRNA transcript abundance for each analyte not only demonstrated the robust nature of the pix H-score algorithm but also validated the pathologist scores.

There are two basic approaches to quantifying biomarker expression from histology images. One approach utilizes cell segmentation and quantifies markers per unit cell whereas a second approach avoids cell segmentation and quantifies markers per unit pixel. In this manuscript, we compared both approaches to quantify biomarker levels from immunohistochemistry images. Unlike the H-score and the Allred score, the pix H-score is a pixel-based algorithm that does not rely on the identification of individual cells and their subcellular compartments. This reduces the computational complexity of the pix H-score and renders its implementation in two different software packages as relatively straightforward.

In our case, the IHC assay for each biomarker was carried out using a different brand of instrument (PD-L1 – Ventana, P-cadherin – DAKO, and 5T4 – Leica Bond RX). Similarly, the slides were scanned using different whole-slide scanners (PD-L1 and P-cadherin - different Aperio AT2 scanners, and 5T4 – Hamamatsu NanoZoomer). These differences could introduce variations in the colorimetric composition of the IHC images that can impact downstream image analysis. Our observation that the Visiopharm and the HALO versions of pix H-score exhibited similar performance suggests that the pix H-score is a robust algorithm for estimating IHC biomarker abundance in whole-slide images. This is especially relevant due to the proprietary nature of these software packages which precludes users from understanding several technical aspects of the image analysis workflow. For instance, the specific details regarding the color deconvolution algorithm, which is a key pre-processing step, are not accessible to the user in either Visiopharm or HALO. Consequently, while implementing the pix H-score we did not know how similar the output of the color deconvolution step (i.e. hematoxylin and DAB channels) would be in the two software packages.

An important question arises as to why the DIA based H-score exhibited very different performance for P-cadherin but not for PD-L1. The H-score algorithm relied on the detection of individual cells and their subcellular compartments to quantify biomarker levels. Although this task may seem relatively straightforward for a human observer, nucleus/cell-membrane detection and segmentation are challenging image processing problems especially when applied to whole-slide image analysis where there can be considerable variability in the intensity and the sub-cellular localization pattern of the biomarker of interest [21, 22]. In our case, the latter could be a contributing factor since in the P-cadherin IHC images the biomarker signal was localized to both the cell membrane and cytoplasm whereas in the PD-L1 IHC images the biomarker signal was predominantly localized to the cell membrane. Consequently, this may partly explain the reason why for P-cadherin the performance of the DIA H-score was consistently lower than that of the pix H-score whereas for PD-L1 the performance of the DIA H-score was comparable to that of the pix H-score. Not surprisingly others have also reported similar challenges in automated analysis of membrane-localized biomarker signal [23]. This may also partly explain our observation for 5T4 where the correlation between QuPath H-score and 5T4 transcript was lower than the correlation between pix H-score and 5T4 transcript. More specifically, while 5T4 immunoreactivity is predominantly membranous, there is still detectable cytoplasmic signal in the tumor cells which can affect the quantification of the DIA based H-score.

A similar question also arises for the ATM score which, unlike the H-score, is a pixel-based algorithm but also exhibited very different performance for P-cadherin but not for PD-L1 and 5T4. By definition. the ATM score is proportional to the average intensity of the biomarker in the DAB channel. This is calculated by taking all pixels in the DAB channel including pixels that are negative for the biomarker. When the averaging is performed on a whole-slide image, this can significantly dilute the contribution from pixels that are positive for the biomarker resulting in poor performance in predicting biomarker abundance from the IHC image. In contrast, the pix H-score only considers pixels with a valid biomarker signal as DAB positive pixels (based on a user defined threshold). As a result, the pix H-score can robustly estimate biomarker abundance the IHC image. This difference also explains in part the reason for the limited range of values taken by the ATM score when compared to the pix H-score. Specifically, the ATM score for P-cadherin, PD-L1, and 5T4 took values in the range of 24 to 77, 8 to 33, and 11 to 49, respectively. In contrast the pix H-score for P-cadherin, PD-L1, and 5T4 took values in the range of 20 to 207, 1 to 131, and 3 to 170, respectively. The latter values are more comparable to the pathologist H-score, which for P-cadherin, PD-L1, and 5T4 ranged from 17 to 298, 0 to 225, and 0 to 224, respectively.

The application of deep learning methodology for nucleus and cell membrane segmentation holds significant promise as it has been shown to have improved performance over traditional algorithms [24]. However, deep learning methods are supervised approaches that require a substantial amount of training data and extensive validation. In many practical applications, generating such large training datasets is not feasible and algorithm validation can be time consuming. In this regard, the pix H-score algorithm introduced here provides a simple yet robust strategy to quantify biomarker expression even from small datasets, as demonstrated here, and can be implemented within a very short timeframe. An interesting follow up study would be to compare the performance of the pix H-score algorithm with deep learning based, scoring approaches.

We note that while our results are encouraging and show the potential for the pix H-score in scoring membrane biomarkers, the algorithm can benefit from additional validation for other biomarkers. Also, the effect of pre-analytical variables (e.g., cold ischemia time, age of unstained cut slides, etc.) on the performance of the pix H-score needs to be investigated. In addition, the effect of stain variation needs to be explored, which is known to be a notable source of variability in histopathology data. In our current work stain normalization was not necessary, likely due to the small batch size of our datasets which did not exhibit significant colorimetric variability. Although not shown here, we expect the pix H-score to also be applicable to immunofluorescence images. In conclusion, we anticipate the pix H-score to be a useful addition to the digital image analysis toolbox for a fast, reproducible and objective strategy to quantify biomarker expression from immunolabeled tissue sections.

## ACKNOWLEDGEMENTS

We thank Shawn O’Neil and Timothy Affolter for critical reading of the manuscript.

## REFERENCES

1. Aeffner, F., et al., Introduction to Digital Image Analysis in Whole-slide Imaging: A White Paper from the Digital Pathology Association. J Pathol Inform, 2019 10: p. 9.

2. Meyerholz, D.K. and A.P. Beck, Principles and approaches for reproducible scoring of tissue stains in research. Lab Invest, 2018. 98(7): p. 844–855.

3. Aeffner, F., et al., Commentary: Roles for Pathologists in a High-throughput Image Analysis Team. Toxicol Pathol, 2016. 44(6): p. 825–34.

4. Brunnstrom, H., et al., PD-L1 immunohistochemistry in clinical diagnostics of lung cancer: inter-pathologist variability is higher than assay variability. Mod Pathol, 2017. 30(10): p. 1411–1421.

5. Gomes, D.S., et al., Inter-observer variability between general pathologists and a specialist in breast pathology in the diagnosis of lobular neoplasia, columnar cell lesions, atypical ductal hyperplasia and ductal carcinoma in situ of the breast. Diagn Pathol, 2014. 9: p. 121.

6. Hirsch, F.R., et al., PD-L1 Immunohistochemistry Assays for Lung Cancer: Results from Phase 1 of the Blueprint PD-L1 IHC Assay Comparison Project. J Thorac Oncol, 2017. 12(2): p. 208–222.

7. Rimm, D.L., et al., A Prospective, Multi-institutional, Pathologist-Based Assessment of 4 Immunohistochemistry Assays for PD-L1 Expression in Non-Small Cell Lung Cancer. JAMA Oncol, 2017. 3(8): p. 1051–1058.

8. Rizzardi, A.E., et al., Quantitative comparison and reproducibility of pathologist scoring and digital image analysis of estrogen receptor beta2 immunohistochemistry in prostate cancer. Diagn Pathol, 2016. 11(1): p. 63.

9. Barnes, M., et al., Whole tumor section quantitative image analysis maximizes between-pathologists’ reproducibility for clinical immunohistochemistry-based biomarkers. Lab Invest, 2017. 97(12): p. 1508–1515.

10. Tsao, M.S., et al., PD-L1 Immunohistochemistry Comparability Study in Real-Life Clinical Samples: Results of Blueprint Phase 2 Project. J Thorac Oncol, 2018. 13(9): p. 1302–1311.

11. Stalhammar, G., et al., Digital image analysis outperforms manual biomarker assessment in breast cancer. Mod Pathol, 2016. 29(4): p. 318–29.

12. Aeffner, F., et al., The Gold Standard Paradox in Digital Image Analysis: Manual Versus Automated Scoring as Ground Truth. Arch Pathol Lab Med, 2017. 141(9): p. 1267–1275.

13. Choudhury, K.R., et al., A robust automated measure of average antibody staining in immunohistochemistry images. J Histochem Cytochem, 2010. 58(2): p. 95–107.

14. Camp, R.L., G.G. Chung, and D.L. Rimm, Automated subcellular localization and quantification of protein expression in tissue microarrays. Nat Med, 2002. 8(11): p. 1323–7.

15. Pirie-Shepherd, S.R., et al., Detecting expression of 5T4 in CTCs and tumor samples from NSCLC patients. PLoS One, 2017. 12(7): p. e0179561.

16. Bankhead, P., et al., QuPath: Open source software for digital pathology image analysis. Sci Rep, 2017. 7(1): p. 16878.

17. Steiger, J.H., Tests for comparing elements of a correlation matrix. Psychological Bulletin, 1980. 87(2): p. 245–251.

18. Williams, E.J., The Comparison of Regression Variables. Journal of the Royal Statistical Society: Series B (Methodological), 1959. 21(2): p. 396–399.

19. Kovacs, A., J. Dhillon, and R.A. Walker, Expression of P-cadherin, but not E-cadherin or N-cadherin, relates to pathological and functional differentiation of breast carcinomas. Mol Pathol, 2003. 56(6): p. 318–22.

20. Paredes, J., et al., P-cadherin overexpression is an indicator of clinical outcome in invasive breast carcinomas and is associated with CDH3 promoter hypomethylation. Clin Cancer Res, 2005. 11(16): p. 5869–77.

21. Irshad, H., et al., Methods for nuclei detection, segmentation, and classification in digital histopathology: a review-current status and future potential. IEEE Rev Biomed Eng, 2014. 7: p. 97–114.

22. Xing, F. and L. Yang, Robust Nucleus/Cell Detection and Segmentation in Digital Pathology and Microscopy Images: A Comprehensive Review. IEEE Rev Biomed Eng, 2016. 9: p. 234–63.

23. Lopes, N., et al., Digital image analysis of multiplex fluorescence IHC in colorectal cancer recognizes the prognostic value of CDX2 and its negative correlation with SOX2. Lab Invest, 2020. 100(1): p. 120–134.

24. Caicedo, J.C., et al., Evaluation of Deep Learning Strategies for Nucleus Segmentation in Fluorescence Images. Cytometry A, 2019. 95(9): p. 952–965.

